# Model-based detection and analysis of introgressed Neanderthal ancestry in modern humans

**DOI:** 10.1101/227660

**Authors:** Matthias Steinrücken, Jeffrey P. Spence, John A. Kamm, Emilia Wieczorek, Yun S. Song

## Abstract

Genetic evidence has revealed that the ancestors of modern human populations outside of Africa and their hominin sister groups, notably the Neanderthals, exchanged genetic material in the past. The distribution of these introgressed sequence-tracts along modern-day human genomes provides insight into the ancient structure and migration patterns of these archaic populations. Furthermore, it facilitates studying the selective processes that lead to the accumulation or depletion of introgressed genetic variation. Recent studies have developed methods to localize these introgressed regions, reporting long regions that are depleted of Neanderthal introgression and enriched in genes, suggesting negative selection against the Neanderthal variants. On the other hand, enriched Neanderthal ancestry in hair- and skin-related genes suggests that some introgressed variants facilitated adaptation to new environments. Here, we present a model-based method called `diCal-admix` and apply it to detect tracts of Neanderthal introgression in modern humans. We demonstrate its efficiency and accuracy through extensive simulations. We use our method to detect introgressed regions in modern human individuals from the 1000 Genomes Project, using a high coverage genome from a Neanderthal individual from the Altai mountains as reference. Our introgression detection results and findings concerning their functional implications are largely concordant with previous studies, and are consistent with weak selection against Neanderthal ancestry. We find some evidence that selection against Neanderthal ancestry was due to higher genetic load in Neanderthals, resulting from small effective population size, rather than Dobzhansky-Müller incompatibilities. Finally, we investigate the role of the X-chromosome in the divergence between Neanderthals and modern humans.

## 1 Introduction

In recent years, researchers have gathered an increasing amount of high-quality genomic sequencing data from human individuals that lived thousands of years ago (Mathieson et al., 2015) and individuals of extinct hominin sister groups (Prüfer et al., 2014; Meyer et al., 2012). These ancient samples provide unprecedented opportunities to elucidate the evolution of modern human populations and their relation to other hominins. Previous genetic evidence revealed that the ancestors of non-African humans exchanged genetic material with Neanderthal individuals after emerging out of Africa. Traces of this introgression can still be found in the genomes of modern-day humans. The emerging high-quality genomic sequence data for ancient hominins not only confirms these findings, but allows detection of the exact location of these introgressed sequence fragments in the genomes of modern human individuals. Uncovering these tracts has received much attention in recent years (Sankararaman et al., 2014, 2016; Vernot and Akey, 2014; Vernot et al., 2016).

To detect these tracts of Neanderthal introgression, Sankararaman et al. (2014) developed a machinelearning based approach, that operates on suitably chosen “features” of the genetic data. The authors recently extended this approach to jointly detect Neanderthal and Denisovan introgression (Sankararaman et al., 2016); the latter is found to be more prevalent in Oceania and Southeast-Asia. Addressing the same question, Vernot and Akey (2014) developed a different approach to detect tracts of Neanderthal introgression, based on sequence identity and divergence, which the authors also extended to the joint detection of Neanderthal and Denisovan introgression (Vernot et al., 2016). These studies report long regions in modern non-African individuals that are depleted of Neanderthal ancestry and enriched in genes, suggesting general negative selection against Neanderthal variants in genes. Furthermore, they report some allelic variants associated with genetic diseases in genome-wide association studies (GWAS) that might have originated from the Neanderthal population. Yet, the studies report enriched Neanderthal ancestry in hair and skin related genes (keratin pathways), which suggests that these introgressed variants could have helped modern non-African populations to adapt to their local environments. In addition to these findings, the availability of the Neanderthal introgression maps in modern humans has led to numerous follow-up studies, investigating the various functional, evolutionary and medical implications of archaic hominin introgression into modern humans (Dannemann et al., 2017; Gittelman et al., 2016; Rogers, 2015; Simonti et al., 2016; Schumer et al., 2017).

In this article, we present a modification of the method `diCal 2.0`, previously developed by Steinrücken et al. (2015) for the inference of complex demographic histories that we call `diCal-admix`. This modification can be used to efficiently detect tracts of introgressed Neanderthal DNA. It is based on a hidden Markov model (HMM) approach that explicitly accounts for the underlying demographic history relating modern human and Neanderthal populations, including the introgression event.

We first present our model-based method for detecting Neanderthal introgression and demonstrate through extensive simulations that our method is able to efficiently and accurately detect introgression in simulated data. We then apply our method to sequence data of modern humans from the 1000 Genomes Project and a high coverage genome from a Neanderthal individual from the Altai mountains (Prüfer et al., 2014). Our results are in general agreement with previously obtained results, and we discuss similarities, differences, and their functional implications, especially with respect to Dobzhansky-Müller incompatibilities suggested by Sankararaman et al. (2014).

## 2 Materials and Methods

### 2.1 Overview of our method

The method to detect Neanderthal introgression that we present here accounts explicitly for the underlying demographic history relating modern humans and Neanderthals. Therefore we briefly present some of the key features of this demographic model. Researchers have studied various aspects of the ancestral relations between modern African and non-African individuals using different methodologies. Furthermore, a large number of studies have investigated the divergence of Neanderthals and modern humans. These studies resulted in several different, albeit largely consistent estimates for the relevant demographic parameters, and we follow Sankararaman et al. (2014) (specifically Figure SI2.1 of that paper) here for consistency.

The demographic model is depicted in Figure 1(a). The size of the ancestral population, the size of the population ancestral to modern humans, and the size of the African population are set to *N* = 13,000, which is consistent with the estimates provided by Gutenkunst et al. (2009). The size of the non-African population after the split is set to *N* = 2, 000. The size of the Neanderthal population is set to *N* = 2, 000 as well, since it has been shown by Prüfer et al. (2014) that the Neanderthal population size declined rapidly as the population neared extinction. The more recent small size will have a stronger impact on genetic variation than the larger ancestral size. Several studies (Gutenkunst et al., 2009) report a strong population bottleneck in European and Asian populations after the out-of-Africa event, followed by rapid exponential population growth, and Sankararaman et al. (2014) incorporate this into their demographic model. As we will detail later, our method considers each non-African haplotype one at a time. The genetic processes along a single ancestral lineage are not affected by the exact population size history, and thus we do not explicitly include these details in the demographic model used here.

**Figure 1:**
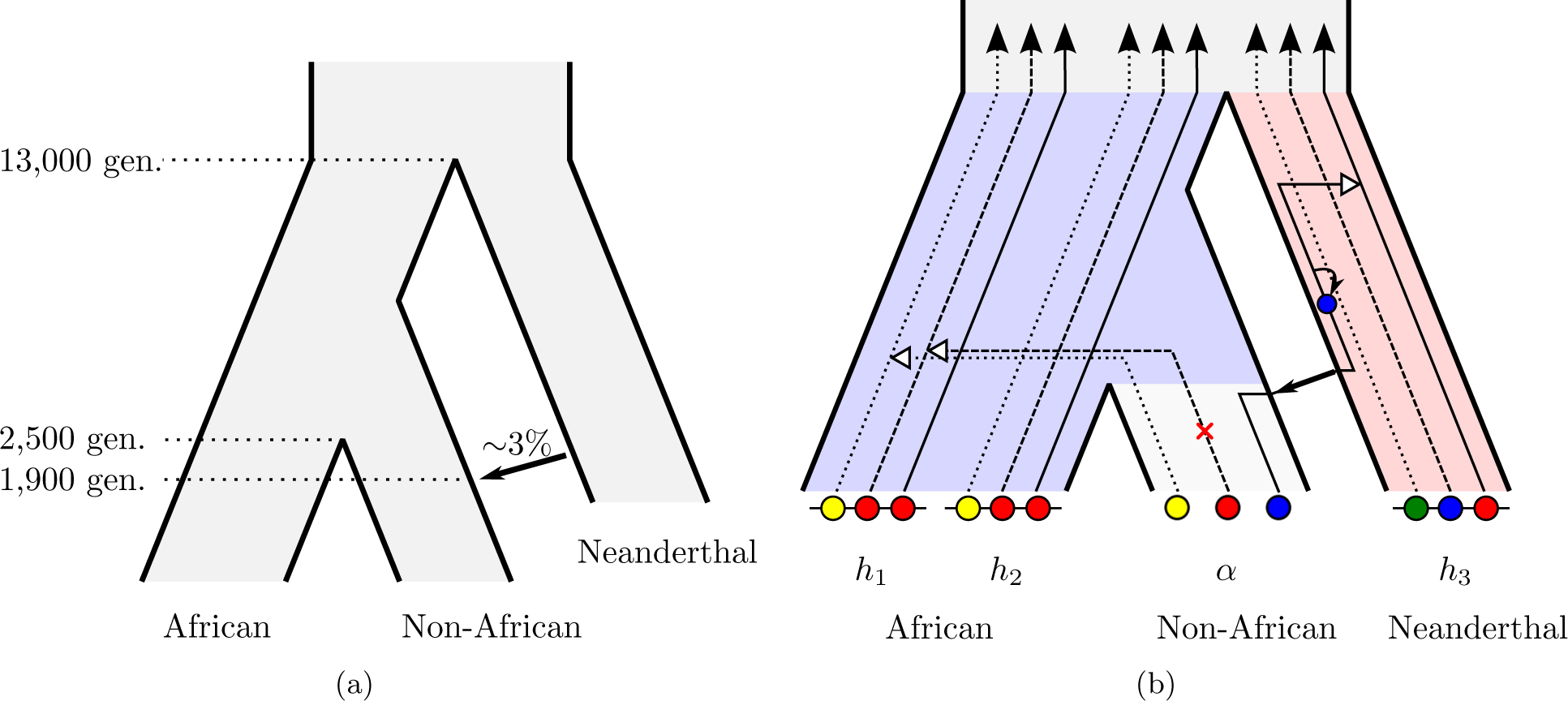
Illustration of the demographic model and the coalescent hidden Markov model. (a) A sketch of the demographic history relating African, non-African, and the Neanderthal population that contains the key features. (b) The Conditional Sampling Distribution applied to the detection of Neanderthal introgression. At each locus, the ancestral lineage of the focal Non-African (*α*) haplotype coalesces either with an African lineage (*h*_1_ or *h*_2_), or a lineage in the Neanderthal population (*h*_3_) through the introgression event.

Like Sankararaman et al. (2014), we set the time of divergence between modern humans and Neanderthals, *T*_nean_, to 13,000 generations ago, which corresponds to 325 kya (Meyer et al., 2012), assuming a generation time of 25 years. Furthermore, the split between African and Non-African *T*_div_ is set to 2,500 generations ago, or 62.5 kya, and the time of the introgression or admixture event *T*_admix_ is set to 1, 900 generations ago (Sankararaman et al., 2012), which corresponds to 47.5 kya (Prüfer et al., 2014). Finally, the introgression coefficient is set as 3%, that is, a non-African individual at the time of introgression had a 3% chance that its parent was a Neanderthal individual. This is consistent with previous estimates of this quantity obtained by Green et al. (2010) and Juric et al. (2016). In Section 2.2, we will demonstrate the robustness of our method to misspecification of key parameters. It has been debated whether all non-African populations received genetic material form the Neanderthals in only one or more than one introgression events. Here we focus on one event, and will defer such investigations to future work.

Before we describe our method to detect tracts of Neanderthal introgression, we provide a brief overview of the methods developed by Sankararaman et al. (2014) and Vernot and Akey (2014) for comparison. In Sankararaman et al. (2014), the authors employ a machine learning approached based on Conditional Random Fields, discriminative analogs of HMMs. To apply this framework, the authors represent the genotype data of the reference Neanderthal, the reference African population, and the focal non-African population in terms of “features” that are informative to distinguish between introgressed and non-introgressed sequence tracts. The authors chose three different classes of features: the distribution of alleles at informative SNPs; a measure of sequence divergence between the focal individual and both reference populations; and a feature to match the length distribution of the observed tracts to be consistent with the expectation from an introgression event 37–86 kya. The authors train the model on data simulated under a demographic scenario similar to Figure 1(a), and apply the trained model to detect introgression tracts in individuals from the 1000 Genomes dataset (The 1000 Genomes Project Consortium, 2012). A modified version of this methodology was applied in Sankararaman et al. (2016) to detect Denisovan ancestry in Southeast-Asians.

Vernot and Akey (2014) developed a two-stage procedure to detect introgression tracts. In the first stage, the authors computed *S*\* statistics (Plagnol and Wall, 2006) of the genomic data in sliding windows. This statistic is sensitive to increased levels of diversity in high linkage disequilibrium, indicating a more ancient *T*_MRCA_ of a given region, and also considers the tract length to detect archaic introgression. Notably, this stage does not require a reference sequence for an archaic individual. In the second stage, the authors then proceed to compare the identified segments to the reference Neanderthal genome, to reliably identify Neanderthal introgression. The same two-stage approach was applied by Vernot et al. (2016) to identify Denisovan introgression.

Here, we apply a modified version of the method `diCal 2.0`, developed by Steinrücken et al. (2015) for inference of ancient demographies, to detect sequence tracts of Neanderthal DNA introgressed into modern humans. The method is based on the conditional sampling distribution (CSD) (Paul et al., 2011; Paul and Song, 2010; Steinrücken et al., 2013), which is similar to the copying model of Li and Stephens (2003), and is depicted in Figure 1(b). The CSD describes the distribution of sampling an additional focal genome or haplotype, conditional on having already observed a certain set of haplotypes. Steinrücken et al. (2015) introduce a version of the CSD that can be applied to haplotypes sampled from several subpopulations, accounting explicitly for the underlying demographic history. Under this model, the unknown genealogy relating the already observed haplotypes is approximated by a trunk genealogy of unchanging ancestral lineages extending infinitely into the past. At each locus, the ancestral lineage of the additional haplotype is absorbed into a lineage of the trunk. The dynamics of absorption depends on the underlying demographic history. In brief, lineages in different subpopulations cannot coalesce, unless continuous or point migration is possible at given rates, and the likelihood of coalescence is larger in small populations, but decreases in large populations. If an ancestral recombination event separates two loci, the haplotype of absorption and the time of absorption can change, thus different genomic segments can be copied from different haplotypes in the trunk, and the additional haplotype is realized as a mosaic of the observed haplotypes. The CSD can be implemented as an HMM along the genome of the additional haplotype, where the hidden state is the trunk haplotype that the genetic material is currently copied from, and a time of absorption. This absorption time is proportional to the likelihood that mutations can alter the genetic type at a given locus. Steinrücken et al. (2015) derived the emission and transition probabilities for the underlying HMM under general demographic models.

This CSD can be applied to detect introgressed tracts of Neanderthal ancestry in modern humans as follows. First, fix the underlying demography given in Figure 1(a) that relates modern African populations, modern non-African populations, and the Neanderthal population. The introgression event is modeled as a point migration. As depicted in Figure 1(b), the haplotypes sampled in the African population and a Neanderthal haplotype are used as the trunk haplotypes in their respective sub-population. Then, each non-African sample is, in turn, used as the additional haplotype in the non-African sub-population. Computing the forward and backward algorithm under the HMM for this CSD yields a marginal posterior distribution over the hidden states at each locus. Recall that these hidden states consist of both an absorption time and an absorbing haplotype. Marginalizing over the absorption time results in a posterior distribution over the trunk haplotypes, and, grouping haplotypes by sub-population, this gives a probability at each locus that the locus in the non-African haplotype is obtained from either an African (modern human) ancestor, or from an Neanderthal ancestor through introgression. Note that using an explicit time for the introgression event in the demographic model implicitly specifies a prior distribution for the length of the introgressed tracts. The software implementation of this method, `diCal-admix`, is available at http://dical-admix.sourceforge.net/.

### 2.2 Simulation study

To demonstrate that `diCal-admix` can be used to accurately and efficiently identify tracts of Neanderthal introgression, we performed an extensive simulation study. To this end, we used the coalescent simulator `msprime` (Kelleher et al., 2016) to simulate sequence data under the demographic model given in Figure 1(a). Specifically, we simulated 176 haplotypes in the African population, one haplotype in the Neanderthal population, and 20 in the focal non-African population. For each such dataset we simulated 20 Mbp of sequence data, using a per generation population scaled mutation and recombination rate of 0.0005. We simulated 50 replicates in each scenario, and estimated the introgression tracts on each focal haplotype.

To investigate how robust our method is to misspecification of the demographic model used for inference, our simulation study was two-fold. First, we simulated data under the demographic model given in Figure 1(a), and varied the demographic model used for the analysis. We kept all parameters fixed and only varied one focal parameter at a time. We varied the divergence time between modern humans and Neanderthal, using 0.5 and 2 times *T*_nean_ = 13, 000 generations, and the divergence time between Africans and non-Africans, using 0.8 and 2 times *T*_div_ = 2, 500 generations. Moreover, we varied the fraction of introgressed Neanderthal individuals, using 0.5 and 2 times `admix` = 3%, and the time of the introgression event, using 0.8 and 1.25 times *T*_admix_ = 1, 900 generations.

For simulated data, the true introgression status at each locus is known. Running `diCal-admix` to detect introgressed tracts in the simulated non-African individuals yields a posterior probability of introgression at each locus. Using different thresholds on this posterior probabilities for calling a locus introgressed, we can assess the true and false positive rate and the precision to generate receiver operating characteristic (ROC) and precision-recall curves for each analysis. Figure 2(a) shows these curves when the data simulated under the “true” model (Figure 1(a)) are analyzed using the different demographic models. We observe an overall good performance and robustness against misspecification of the parameters for analyzing the data. From these plots, we determined that a threshold of 0.42 yields a good balance of the different performance metrics. We indicated this threshold along the curves by a red cross. Note that it is possible to increase the true positive rate by lowering the threshold, however, the decrease in precision would be more severe. Thus, we used this threshold in the remainder to call introgression tracts in the 1000 Genomes dataset.

**Figure 2:**
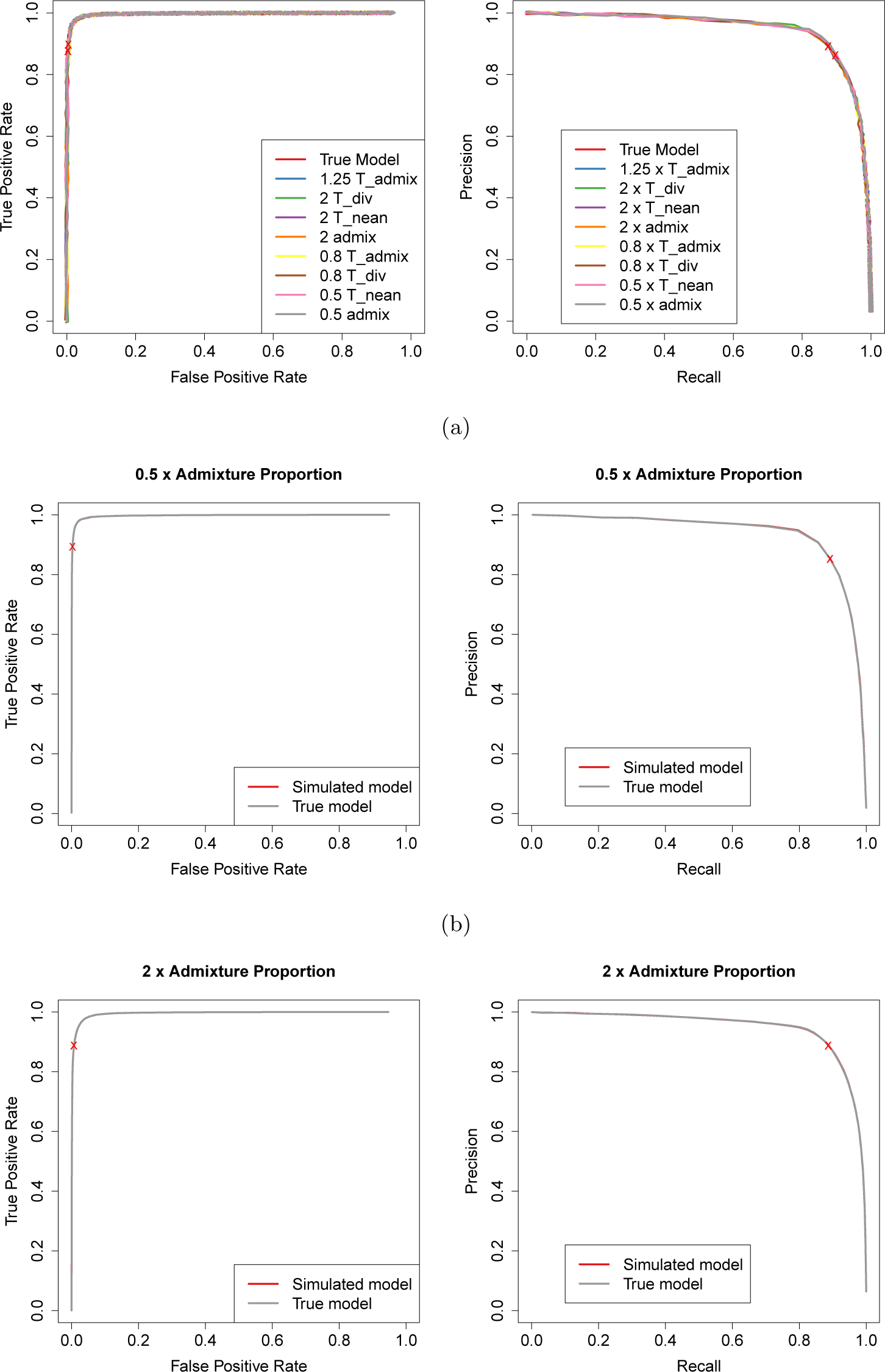
Receiver operating characteristic and Precision-recall curves from the simulation study. (a) For data simulated under the “true” model, and the parameters for the analysis varied. We added a small amount of random “jitter” to the curves since they are hard to distinguish otherwise. (b) For data simulated under a model where the admixture proportion is 1.5% and analyzed using the same model and the “true” model. (c) For data simulated under a model where the admixture proportion is 6% and analyzed using the same model and the “true” model.

For the second part of the simulation study, we simulated data under the different demographic models. We then analyzed each dataset using the same parameters as used for the simulation on the one hand, and using the parameters of the “true” model (Figure 1(a)) on the other hand. The ROC and the Precision-recall curves for varying the introgression percentage are depicted in Figure 2(b) and Figure 2(c), and the curves for the remaining scenarios are given in the Supporting Information SI.2. Again, we observe that misspecifying the parameters of the analysis does not affect the performance substantially. The ROC curves demonstrate a good performance in terms of true and false positive rate in most scenarios, however, the precision-recall curves exhibit a poorer performance in some scenarios. This is to be expected, as in the some scenarios, the Neanderthal and African population are genetically closer, for example, when the divergence time between Africans and non-Africans is increased, or the divergence time between Neanderthal and modern humans is decreased. If the populations are more closely related, it becomes more difficult to distinguish introgressed variation from variation shared between modern human populations.

## 3 Results

### 3.1 Neanderthal introgression in the 1000 Genomes data

We applied `diCal-admix` to detect tracts of Neanderthal introgression in non-African individuals from the Phase I dataset of the 1000 Genomes Project (The 1000 Genomes Project Consortium, 2012), focusing on Europeans (CEU) and East Asians (CHB and CHS) in particular. Specifically, we used the 88 YRI individuals (176 haplotypes) from this dataset as reference African haplotypes, assumed to have no introgressed genetic material from Neanderthals, that serve essentially as a modern human reference panel. We used `diCal-admix` to compute the marginal posterior introgression probability along the genomic sequences of each of the 85 CEU individuals (170 haplotypes), 97 CHB individuals (194 haplotypes), and 100 CHS individuals (200 haplotypes) in turn. We used a high-coverage genomic sequence from an Altai Neanderthal individual (Prüfer et al., 2014) as a Neanderthal reference. Prüfer et al. (2014) presented different genome alignability filters (Prüfer et al., 2014, SI 5b), and we used the `map35_50%`-filter, since this filter was suggested by the authors to be most appropriate for population genomic analyses.

`diCal-admix` requires that the genomic data be phased into haplotype sequences. The 1000 Genomes dataset is computationally phased, so we could use this data as provided; however, the diploid sequence of the Neanderthal individual cannot be phased using standard statistical methods. We instead used an additional pre-processing step to obtain a pseudo-haplotype sequence. As noted by Prüfer et al. (2014), the Altai Neanderthal individual exhibits only a sixth of the heterozygosity of modern non-African individuals. Thus, the number of ambiguous sites that require phasing is small. We tested three different methods to obtain a haplotype allele for these remaining sites: choosing an allele uniformly at random, using the ancestral allele only, and using the derived allele only, where the ancestral states at each locus were determined using a six-primate consensus (Paten et al., 2008). We observed little difference in our results between the different approaches, and thus we only present the results using the ancestral-allele approach. We used a mutation rate of 1.25 × 10^−8^ per site per generation (Scally and Durbin, 2012) and chromosome-specific recombination rates obtained by averaging the fine-scale rates provided by Kong et al. (2010). Note that we made the simplifying assumption that the recombination rate is constant within each chromosome. Due to computational considerations, we did not compute the posterior at every genomic site, but rather grouped sites together into 500 bp windows; the details of this procedure are provided in Steinrucken et al. (2015). Furthermore, we applied a moving average filter of length 50 kbp to the raw posterior in a post-processing step, to smooth sudden changes.

Moreover, we obtained from Sankararaman et al. (2014) the likelihoods of Neanderthal introgression that they computed for the same individuals. To compare these calls obtained at the SNPs to the ones obtained using `diCal-admix`, we interpolated the Sankararaman et al. likelihoods at the position in the middle of the 500 bp windows employed by `diCal-admix`. As advised by Sankararaman et al., we used a threshold of 0.89 to call Neanderthal introgression tracts. Vernot and Akey (2014) also identified tracts of Neanderthal ancestry in the individuals from the 1000 Genomes dataset, excluding the X-chromosome. We downloaded the population summaries from http://akeylab.princeton.edu/downloads.html and compared them to the results obtained with `diCal-admix`, when possible.

We averaged the marginal posterior obtained using `diCal-admix` across each chromosome and across all CEU individuals, and separately across all CHB+CHS individuals. We performed the same averaging for the posterior probabilities obtained by Sankararaman et al. (2014). Figure 3(a) shows the results for each chromosome in the CEU population, while Figure 3(b) shows the results for the CHB+CHS population. We find an average introgression of 1.03% in the CEU autosomes, and 1.25% in the CHB+CHS autosomes, whereas Sankararaman et al. (2014) report 1.10% and 1.31%, respectively. However, the average amount of introgressed material varies, from as low as 0.20% on chromosome 17 in the CEU population, to as high as 2.30% on chromosome 9 in the CHB+CHS population. Compared to the autosomes, Sankararaman et al. (2014) previously reported a smaller amount of introgressed material on the X-chromosome in CEU (0.16%) as well as in CHB+CHS (0.22%). Similarly, we observed a roughly five-fold decrease on the X-chromosome when compared to the autosomes in CEU (0.20%) and CHB+CHS (0.28%). In general our results were in good agreement with Sankararaman et al. (2014), but we detected less introgressed genetic material on the autosomes, and more introgression on the X-chromosome.

**Figure 3:**
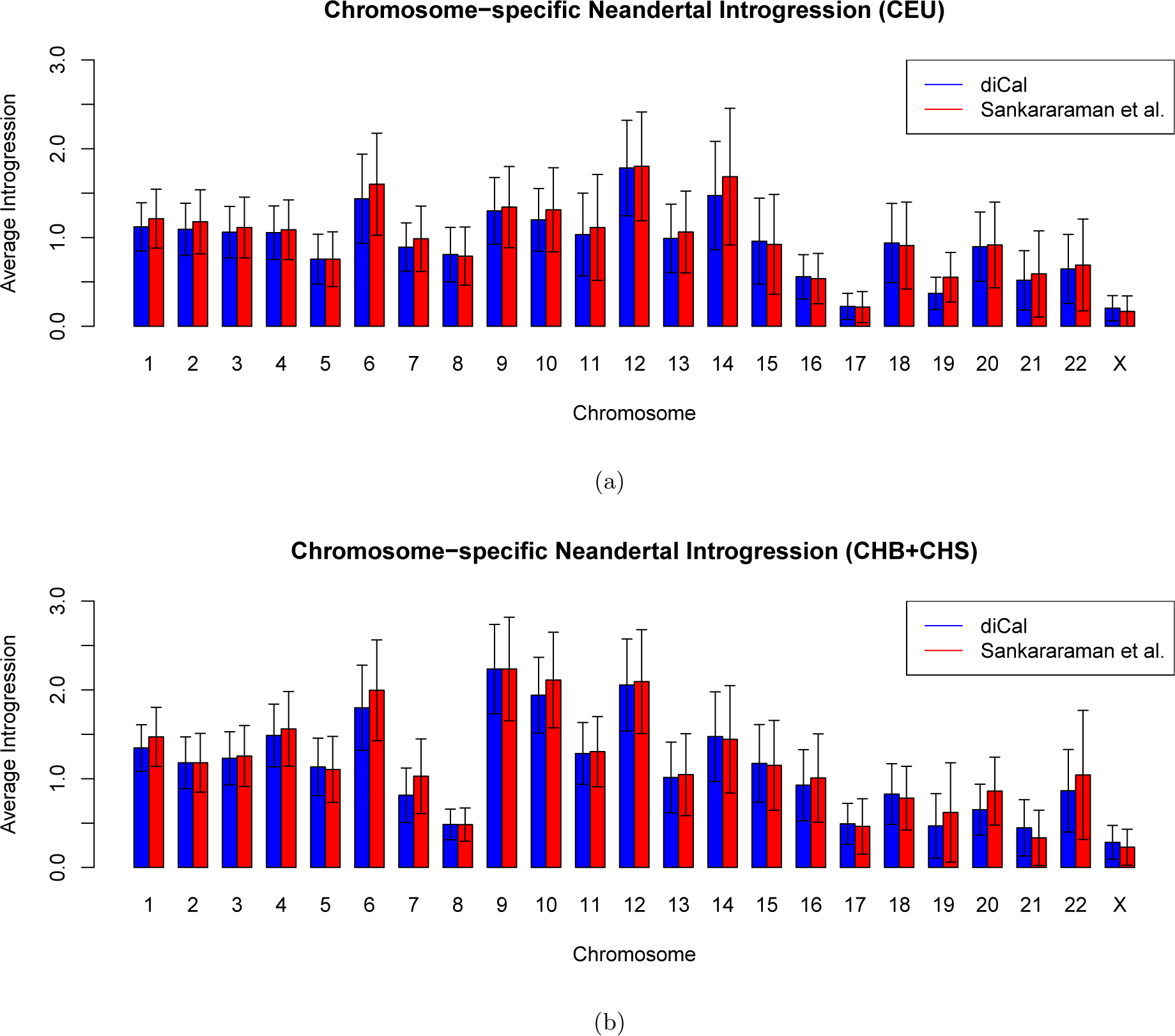
The amount of Neanderthal introgression in modern-day individuals, stratified by chromosome. The height of each bar gives the average introgression, and the whiskers indicate the standard deviation across the sample. (a) CEU. (b) CHB+CHS.

The posterior distributions along the chromosomes allow for a more detailed view of Neanderthal introgression into modern humans as it varies along the genome. We determined whether a given locus is admixed on a particular haplotype by thresholding the posterior generated using `diCal-admix` at 0.42 and thresholding the posterior from Sankararaman et al. (2014) at 0.89. We then averaged these calls across 1 Mbp windows and across the individuals in the respective populations, and plotted the result as piece-wise constant functions. The skyline plots in Figure 4 show the percentage of Neanderthal introgression along chromsome 4 in CEU and the X-chromosome in the CEU population. (In the Supporting Information SI.3, we provide skyline plots for all chromosomes in the CEU and the CHB+CHS population.) In addition, we indicated the regions on the autosomes that were identified in Vernot and Akey (2014) to be introgressed. As mentioned earlier, the X-chromosome was excluded in their study. We see good agreement between the calls made using `diCal-admix` and the calls from Sankararaman et al. (2014). Furthermore, the regions of introgression detected by Vernot and Akey (2014) cluster in regions were the skyline plots indicate introgressed genetic material.

**Figure 4:**
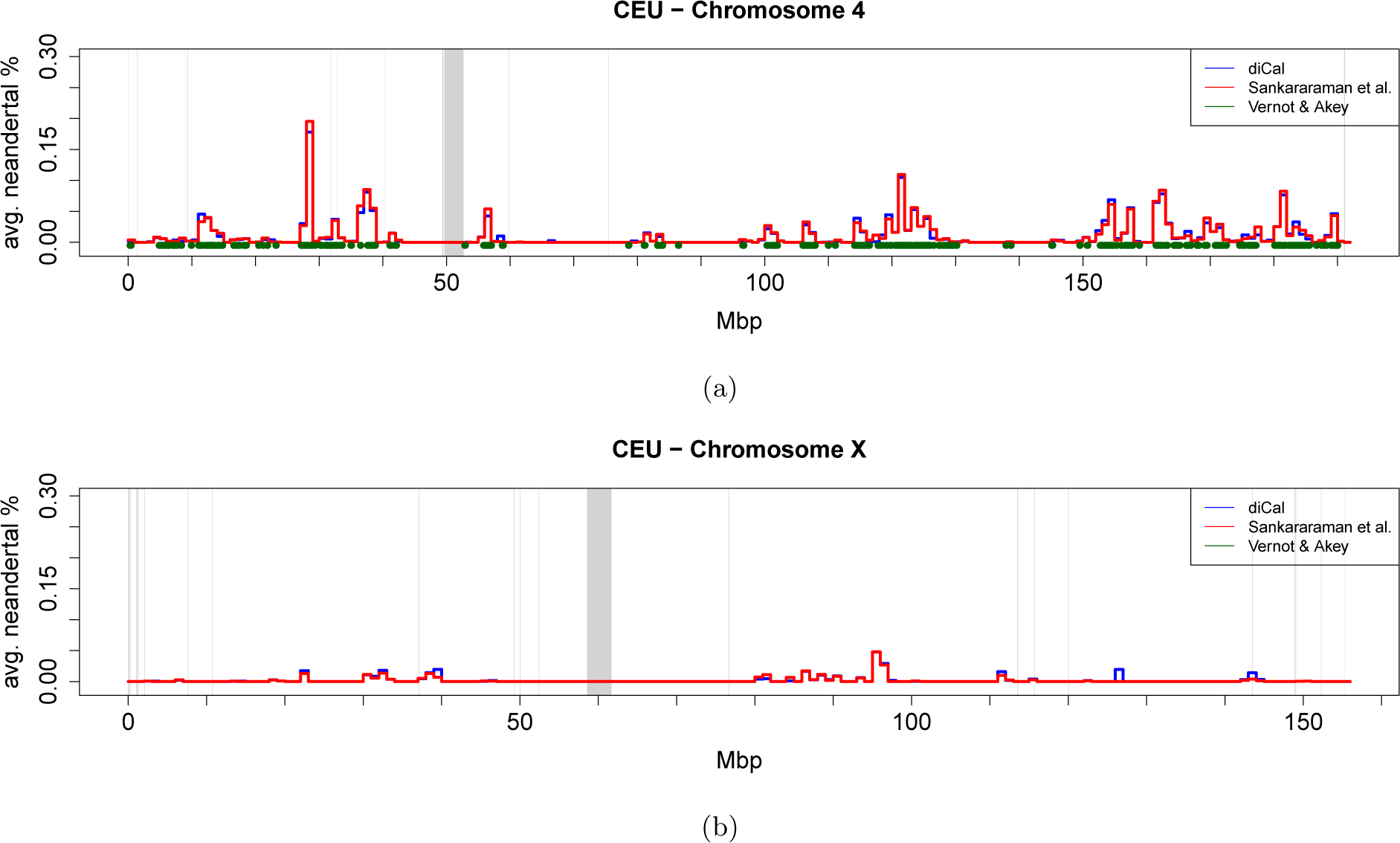
Skyline plot of the amount of Neanderthal introgression in the CEU population, averaged over all individuals in 1 Mbp windows. The results from `diCal-admix` are indicated in blue, and the results from Sankararaman et al. (2014) indicated in red. The regions reported as introgressed by Vernot and Akey (2014) are indicated in green. The gray bars denote the regions were no calls were made in the 1000 Genomes dataset, which include the centromeres. (a) Chromosome 4. (b) Chromosome X.

To investigate the shared features and differences between the introgression call-sets, we generated Venn diagrams. For the `diCal-admix` posterior and the posterior from Sankararaman et al. (2014), we used the aforementioned thresholds to call introgression tracts in the CEU and CHB+CHS individuals. For each individual, we assessed at each locus whether either method, both methods, or no method detected Neanderthal introgression, and averaged these indicators to get population-wide percentages for the whole genome. Figures 5(a) and 5(b) show the Venn diagrams for the different call-sets in the CEU population, for the autosomes and the X-chromosome, respectively. Figures 5(c) and 5(d) depict the results for the CHB+CHS population, on the autosomes and the X-chromosome, respectively. We observe a large overlap between the calls based on `diCal-admix` and Sankararaman et al. (2014) on the autosomes, but less agreement on the X-chromosome. We also generated population-wide introgression maps in each population, called a *tiling path* by Sankararaman et al. (2014). To this end, we identified those regions on the chromosomes where introgression was called for at least one individual in the respective population. We then compared these population-wide introgression maps with the population-level introgression maps published by Vernot and Akey (2014). We generated three-way Venn diagrams for the autosomes in the CEU population and the CHB+CHS population, shown in Figure 6, both in units of percentage of the whole autosome. Again, we observe a large overlap between `diCal-admix` and Sankararaman et al. (2014), but less so with Vernot and Akey (2014). This discordance might be explained to some degree by the fact that in the two-stage procedure of Vernot and Akey (2014) the first step does not use sequence information from the Neanderthals as in the other methods.

**Figure 5:**
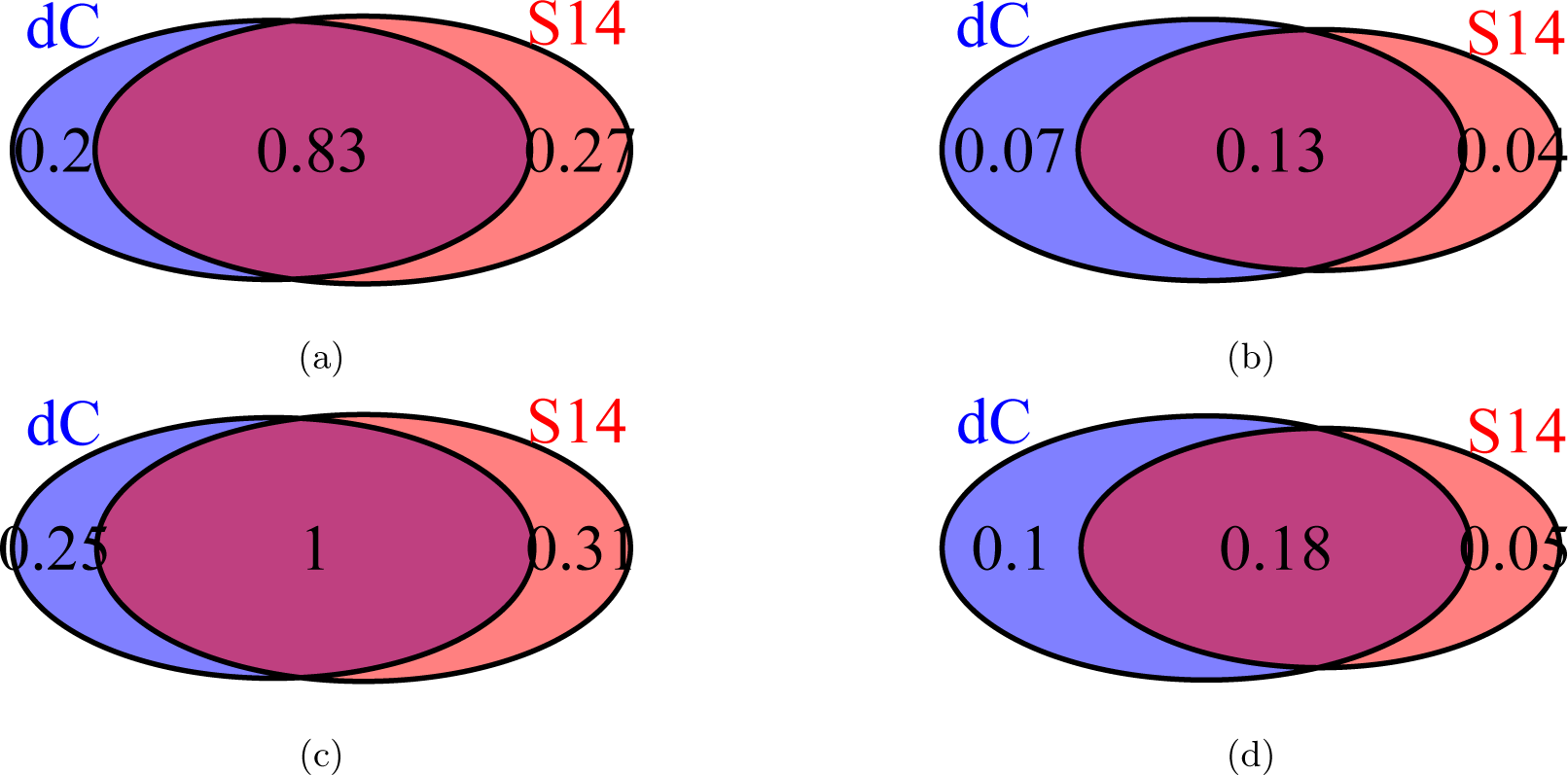
Venn diagrams of the average percentage of introgressed genetic material detected by `diCal-admix` (dC) and by Sankararaman et al. (2014) (S14). (a) For autosomes in the CEU individuals. (b) For the X-chromosome in the CEU individuals. (c) For autosomes in the CHB and CHS individuals. (d) For the X-chromosome in the CHB and CHS individuals.

**Figure 6:**
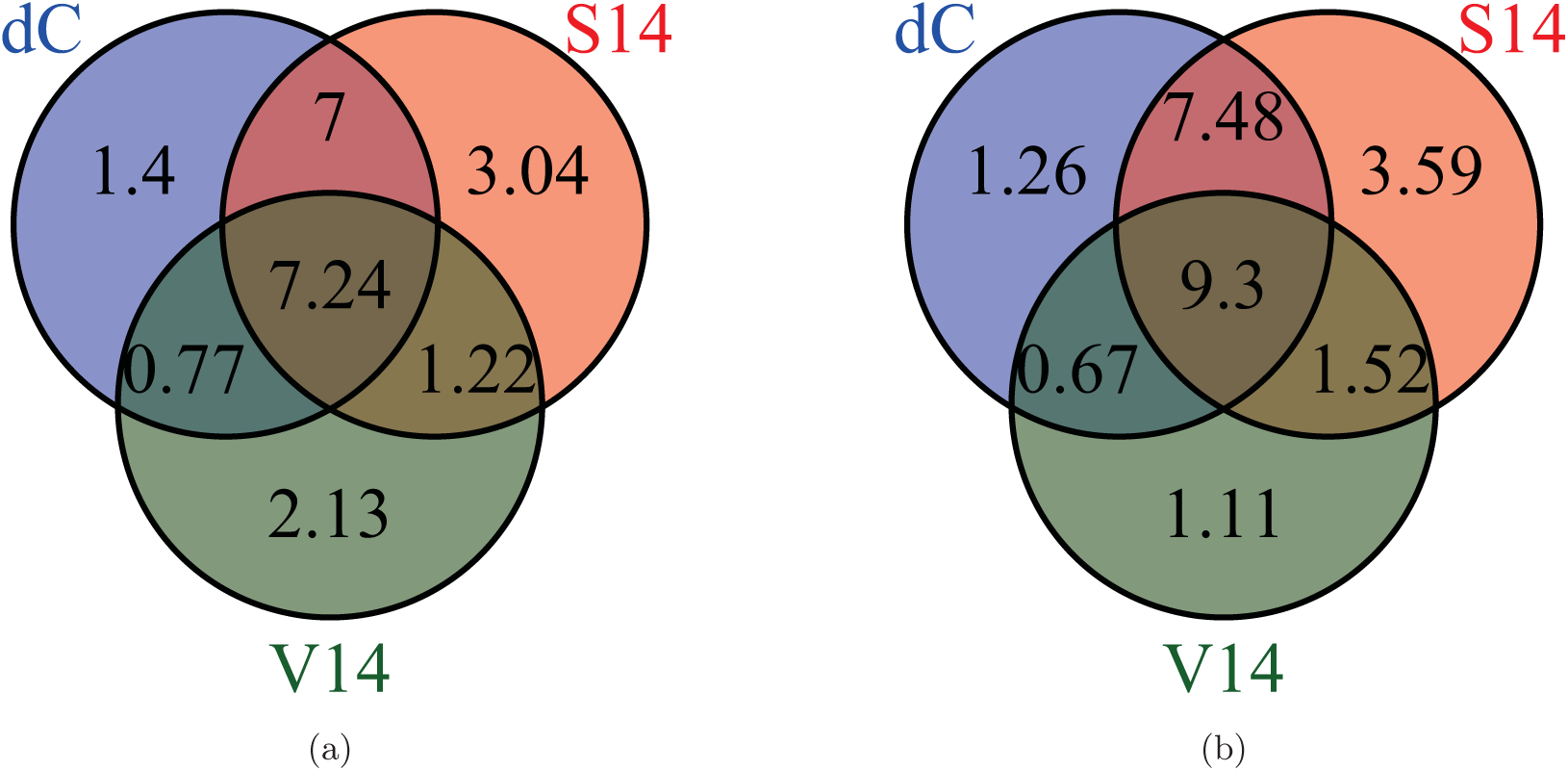
Venn diagrams of the average percentage of regions on the autosomes where Neanderthal introgression was called for at least one individual in the population, by `diCal-admix` (dC), Sankararaman et al. (2014) (S14), and Vernot and Akey (2014) (V14). (a) CEU. (b) CHB+CHS.

We also investigated the distribution of fragment lengths that were detected by the different methods. For all individual from a given population, we counted the number of times an introgression tract of a specific length was detected. Figure 7(a) and 7(c) depict the distributions of the absolute frequencies in the autosomes of the individuals in the CEU population and the CHB+CHS population, respectively. Figure 7(b) and 7(d) show the same distributions for the X-chromosomes. In addition to the empirical tract length distribution obtained from the 1000 Genomes individuals, we plotted the neutral expectation of the absolute frequencies. This neutral expectation of the tract length distribution is computed under the following simple model. Approximating the chromosome as continuous, and considering the introgression tract in an individual at present, the distance between recombination breakpoints is exponentially distributed with parameter *g* × *r*, where r is the per generation per base-pair recombination probability and *g* is the number of generations since the introgression event, because in each generation, there is a chance that recombination breaks down the introgression tract. Here we used *g* = 1, 900 and *r* = 1.19 × 10^−8^ for the autosomes and 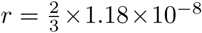 for the X-chromosome. The exponential rate *g* × *r* can also be used to obtain the expected number of sequence tracts in a genome of a certain size, 3% of which are introgressed from an ancestral Neanderthal individual, which yields the expected absolute frequency.

**Figure 7:**
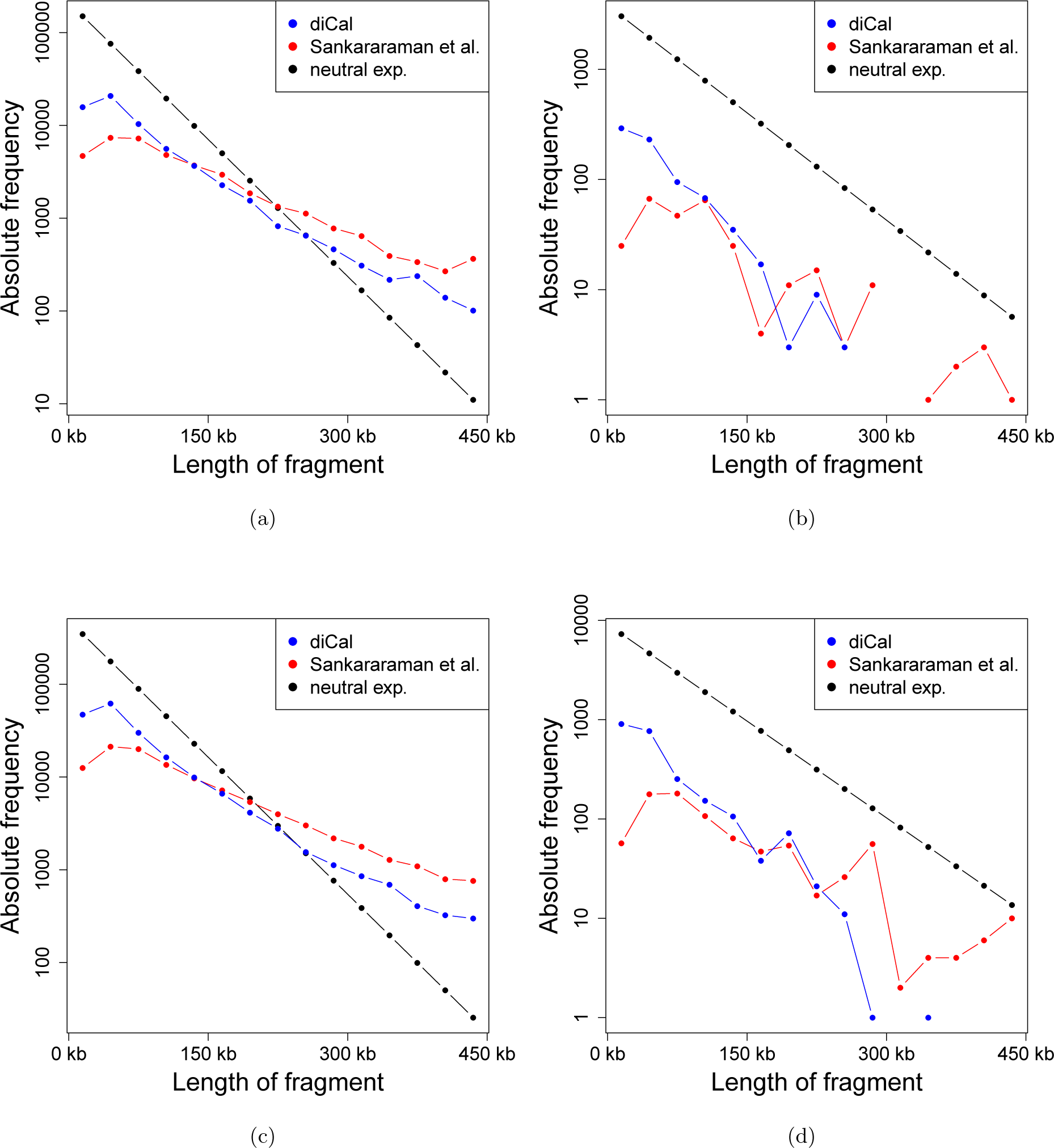
The empirical distribution of the lengths of the introgression tracts, accumulated across individuals. The different length are binned into classes of size 30 kbp. Missing values indicate unobserved classes. (a) Absolute frequency of tracts of a given length on the autosomes in the CEU population. (b) Absolute frequency of tracts of a given length on the X-chromosome in the CEU population. (c) Absolute frequency of tracts of a given length on the autosomes in the CHB+CHS population. (d) Absolute frequency of tracts of a given length on the X-chromosome in the CHB+CHS population.

This simple model for the neutral expectation is certainly oversimplified, but it serves as a first approximation. It is not discernible in these plots whether deviation from the neutral expectation is due to incorrect detection of the tracts, or the true underlying tracts actually being subject to non-neutral evolution. The fact that both methods deviate from the neutral expectation in qualitatively similar ways suggests both factors may be playing a role. However, it is surprising that, for the autosomes, both methods detect more long fragments than expected under the simple neutral model. This would suggest that either there is at least a component of an introgression event that happened more recently than 1, 900 generations ago, or some form of selection is acting that favors longer fragments.

In general, `dical-admix` detects more short fragments and fewer long fragments than reported by Sankararaman et al. (2014). Moreover, the empirical distribution of `dical-admix` is closer to the neutral model. This and the other statistics of the empirical distribution of the introgressed Neanderthal tracts presented in this section suggest that there is merit in applying different methodologies for the detection of introgression. While all methods perform reasonably well on simulated data, they seem to be sensitive to slightly different features of the introgression tracts. Thus we suggest to use the consensus of the three methods for highly confident introgression calls, and use the regions unique to only some of the methods for more exploratory research.

### 3.2 Functional implications of Neanderthal introgression

To explore the functional implications of Neanderthal Introgression we performed a gene ontology (GO) analysis using GOrilla (Eden et al., 2007, 2009), which looks for overrepresentation of GO terms at the top of a ranked list of genes. For each population, we ranked genes by their mean posterior probability of introgression (as determined by the `diCal-admix` posterior decoding) and looked for GO terms associated with either a lack of Neanderthal introgression or an enrichment of introgression. We restricted our analyses to the 500 bp resolution introgression calls where no more than half of the bases are masked by the 1000 Genomes strict mappability mask (The 1000 Genomes Project Consortium, 2012). The results (shown in the Supporting Information SI.1) are broadly concordant between populations. Like Sankararaman et al. (2014), we find that genes associated with hair, skin, and keratin are more likely to be introgressed than other genes, which hints at the possibility of adaptive introgression for these genes. Intriguingly, sensory perception, particularly olfaction, had both genes more likely to be introgressed as well as genes less likely to be introgressed. It is possible that adaptive introgression has played a role for some of these genes, for example by helping to adapt to local environments and that selection has removed introgression at other olfaction-related genes. It is also possible, however, that this is an artifact of either the introgression calls (e.g. due to high amount of polymorphism in ofactory genes Malnic et al. (2004)), or high variance in mean introgression rate (e.g. due to being smaller than other genes, or being spatially clustered in the genome).

We also investigated whether SNPs associated with particular phenotypes are more or less likely to be introgressed on average. To this end, we downloaded the results of 2,419 GWASs that were performed on data from the UK biobank (Sudlow et al., 2015; Global Biobank Engine, 2017) and extracted all of the SNPs that were significant at a genome-wide significance level of 5.0 × 10^−8^ for each GWAS. We then tested whether the mean posterior probability of introgression at these SNPs was significantly higher or lower than expected using our bootstrap-like test (described below). Perhaps due to the large number of tests performed, we did not find any statistically significant results, but a number of tests were nominally significant at the *p* = 0.01 level in both populations and may be of interest for future research. In particular we find that loci associated with vaginal or uterine prolapse are more likely to be introgressed in both populations (nominal *p* = 0.0054 in CEU, nominal *p* = 0.00442 in CHB+CHS), as are loci associated with being treated with the drug cardioplen, which is used to treat heart disease (nominal *p* = 0.0097 in CEU, nominal *p* = 0.0063 in CHB + CHS), and loci associated with being treated with desloratadine, which is used to treat allergies (nominal *p* = 0.0061 in CEU, nominal *p* = 0.0001 in CHB+CHS). We did not find any sets of loci identified by a GWAS that were less likely to be introgressed, perhaps due to the low overall levels of introgression resulting in a lack of power. We again urge caution in interpreting these results, due to the multiple testing burden making the above results not statistically significant.

### 3.3 Causes of selection against Neanderthal introgression

While many recent studies have found evidence that natural selection has acted to remove segments of Neanderthal ancestry (Sankararaman et al., 2014; Harris and Nielsen, 2016; Juric et al., 2016), there is some debate about the precise cause of this selective pressure. Dobzhansky-Müller incompatibilities (DMIs) are a classic explanation for selection acting against introgressed alleles; DMIs are alleles that have arisen separately in each population and are neutral in isolation, but are deleterious when brought together in individuals of hybrid ancestry (Dobzhansky, 1936; Orr, 1995). DMIs have been observed in the hybrids of other species—e.g., *Drosophila simulans* and *D. melanogaster* (Brideau et al., 2006); *Mimulus guttatus* and *M. nasutus* (Fishman and Willis, 2001); and *Ambystoma californiense* and *A. tigrinum mavortium* (Fitzpatrick, 2008)—and were hypothesized (particular in male hybrids) to be the cause of selection against Neanderthal ancestry in modern humans (Sankararaman et al., 2014). The latter arrived at this hypothesis-due to finding significant enrichment of genes expressed in testes in regions of low Neanderthal ancestry. In addition, they observed a substantial reduction of Neanderthal ancestry on the X-chromosome, and it has been observed, in other species, that loci contributing to reduced male fertility in hybrids are concentrated on the X-chromosome (Presgraves, 2008). Our results obtained using `dical-admix` also show a reduction in Neanderthal ancestry on the X-chromosome, but this reduction can be explained without appealing to DMIs, as we will discuss in more detail in Section 3.4. Here, we focus on global features of the genome and how they relate to potential DMIs.

Compared to modern humans and Neanderthals (separated by tens of thousands of generations), however, the species in which DMIs have been observed have been substantially more diverged—e.g., about 20 million generations between *D. simulans* and *D. melanogaster* (Li et al., 1999); 200-500 thousand generations between *M. guttatis* and *M. nasutus* (Brandvain et al., 2014); and 750 thousand to 5 million generations between *A. californiense* and *A. tigrinum mavortium* (Shaffer and McKnight, 1996; Fitzpatrick et al., 2010). Furthermore, the divergence time separating modern humans and Neanderthals is only about a factor of two older than the divergence time between the most diverged human populations (e.g., as inferred by Veeramah et al. (2012) and Gronau et al. (2011) using an updated mutation rate as discussed in Scally and Durbin (2012)) and no DMIs are known to occur in admixtures of modern human populations, raising the question of how such incompatibilities between modern humans and Neanderthals could have arisen so quickly.

Juric et al. (2016) and Harris and Nielsen (2016) recently independently proposed that selection acts against Neanderthal ancestry due to a higher mutational load in Neanderthals rather than DMIs. In their study, Juric et al. developed a likelihood method to explicitly infer the strength of selection against Neanderthal introgression based on the introgression maps obtained by Sankararaman et al. (2014). The authors estimated a selection coefficient for deleterious exonic Neanderthal alleles around –4 × 10^−4^ for the autosomes. Since the estimated coefficient is on the order of the inverse of the effective population size in humans, they hypothesized that the deleterious alleles could have accumulated as a result of the small longterm effective population size in Neanderthals (Prüfer et al., 2014), which reduced the efficacy of selection in this population. When these deleterious alleles entered the larger human population through introgression, they were subjected to more efficient selection, and this led to the observed widespread selection against Neanderthal alleles. The authors use simulations to confirm that the population size history of Neanderthals could have indeed allowed for the accumulation of deleterious alleles on the observed order of magnitude. Harris and Nielsen (2016) arrived at similar conclusions while studying the strength of selection against Neanderthal introgression using forward simulations of autosomal genetic material.

In an attempt to disentangle the DMI and mutational load hypotheses, we performed a number of statistical tests. As detailed below, we found evidence that there was selection against Neanderthal ancestry, but could not find statistically significant evidence that the selection was due to DMIs. We interpret this as evidence in favor of the mutational load hypothesis, although it is possible that more data or more powerful statistical tests may show evidence in favor of the DMI hypothesis.

We repeatedly used a test similar to bootstrapping, which we describe presently and refer to subsequently as the *bootstrap-like test*. When performing hypothesis tests, we must account for the spatial correlation of both our introgression calls and many genomic features of interest (e.g. gene locations or local recombination rates). We also would like to account for uncertainty in the introgression calls themselves. To this end, we left the genomic features of interest in place, and then sampled new introgression calls for each chromosome by drawing, with replacement, non-overlapping 5 Mb segments of our original introgression calls from the same chromosome. We then recalculated our test statistic using this resampled set of introgression calls. Repeating this resampling procedure many times provided an approximate empirical distribution of our test-statistic under the null hypothesis of no association between our introgression calls and the genomic feature of interest. We then used this distribution to compute approximate *p*-values. For all of the tests presented below, we again restricted our analyses to the 500 bp windows where no more than half of the bases were masked by the 1000 Genomes strict mappability mask (The 1000 Genomes Project Consortium, 2012).

To begin, we looked into whether selection against Neanderthal ancestry has occurred. First, note that the admixture proportion has previously been estimated as 3% (Green et al., 2010; Juric et al., 2016). If there was no subsequent “dilution” of Neanderthal ancestry (Sankararaman et al. (2016), but see Vernot and Akey (2015), and see Slatkin and Racimo (2016) for a comprehensive review), then under neutrality we would expect about 3% ancestry on average in present-day populations, and we would expect about half of chromosomes to have more than 3% Neanderthal introgression on average and about half of chromosomes to have less than 3% introgression. Yet, no chromosome in either CEU or CHB+CHS has, on average, more than 2.54% introgression, and most chromosomes have less than 1.5% average Neanderthal ancestry. Thus, under this simple null model we can reject neutrality (*p* = 2.4 × 10^−7^, two-sided sign test, *n* = 23) in both CEU and CHB+CHS. While the above test assumed that the admixture proportion was 3%, we would would be able to reject the null hypothesis of neutrality for any admixture proportion greater than 1.14% in CEU or 1.46% in CHB+CHS, both of which are much lower than the findings of the previous studies discussed above. Indeed, Juric et al. report a confidence a 95% confidence interval of [3.22%, 3.52%] for the admixture proportion in CEU and [3.45%, 3.86%] for the admixture proportion in CHB+CHS (Juric et al., 2016).

To explore if this reduction in Neanderthal ancestry is more pronounced in genic regions, we compared the mean (across individuals and loci) frequency of introgression in regions marked as exons in the RefSeq annotation (O’Leary et al., 2016) to the chromosome-wide mean. The results were largely concordant when we considered transcripts or coding sequences instead of exons, so we present below only results based on exons. For CEU, we found that in 17 out of the 23 chromosomes there is less introgression in genic regions than in the rest of the chromosome, which is statistically significant (*p* = 0.0347, two-sided sign test, *n* = 23). For CHB+CHS, however, we found that only 14 chromosomes have less introgression in genic regions, which is not statistically significant (*p* = 0.405, two-sided sign test, *n* = 23). We also performed our bootstrap-like test to see if there was any significant decrease in genic regions on any chromosome, but we did not find any significant results (unadjusted *p* > 0.05 on all chromosomes). We interpret these results as indicating that either selection against Neanderthal introgression is fairly weak if it is acting on most or all genes, or has only acted on a subset of genes. It is also possible that selection against Neanderthal introgression has acted on genomic elements other than exons, such as regulatory elements.

Meanwhile, we found evidence that a measure of conservation, phastCONS (Siepel et al., 2005), was significantly negatively correlated with mean introgression at a given locus (Spearman’s *ρ* = -0.027, *p* = 0.004 in CEU and *ρ* = -0.023, *p* = 0.031 in CHB+CHS, bootstrap-like test), which indicates that selection was more likely to remove Neanderthal ancestry at highly conserved loci. We also tested if proportion of Neanderthal ancestry was positively correlated with local population-scaled recombination rate (as inferred by Myers et al. (2005)), which would be suggestive of selection against Neanderthal ancestry because regions of high recombination would be more likely to separate neutral regions of Neanderthal ancestry from linked deleterious regions, an idea recently explored elegantly and in more detail by Schumer et al. (2017). Similar to Schumer et al., we found a positive association between local population-scaled recombination rate and frequency of introgression (Spearman’s *ρ* = 0.019, *p* = .029 in CEU and *ρ* = 0.022, *p* = 0.009 in CHB+CHS, bootstrap-like test) lending further credence to the hypothesis that selection is acting against certain regions of Neanderthal ancestry.

If DMIs were the cause of selection against introgression, then we would expect that genes that code for proteins with more binding partners would be less likely to be introgressed; each protein-protein interaction (PPI) can be thought of as a possible DMI. To test this hypothesis, we used the PICKLE2.0 PPI network (Klapa et al., 2013; Gioutlakis et al., 2017) and associated a number of binding partners to each gene by counting the number of PPIs in which the protein coded by that gene participates. We found an insignificant correlation between number of binding partners and mean frequency of introgression in both populations (Spearman’s *ρ* = -0.002, *p* = 0.915 in CEU, *ρ* = 0.007, *p* = 0.648 in CHB+CHS, two-sided bootstrap-like test), which provides indirect evidence against the DMI hypothesis.

As a more direct test of the DMI hypothesis, we tested whether proteins that interact (according to the PICKLE2.0 PPI network) are more likely to be co-introgressed. In particular, for each gene in the PPI network, we say that that gene is introgressed if any part of any of its exons is in a called introgression tract. For each individual we then assign a weight to each edge in the PPI network as follows. Let gene A and gene B be the genes that code for the proteins involved in the interaction corresponding to the edge of interest. If each copy of gene A and gene B (i.e. on autosomes, we assume there are two copies of each gene and on the X-chromosome males have only one copy) has the same ancestry, the edge is assigned a weight of one – in this individual, this interaction is always between proteins from genes of the same ancestry. Meanwhile, if all of the copies of gene A are of one type of ancestry and all of the copies of gene B are of the other type of ancestry, then the edge is assigned a weight of zero – this interaction is never between proteins of the same ancestry. Finally, if either gene has mixed ancestry (i.e. one copy from one ancestry and the other copy from the other ancestry) the edge is assigned a weight of 0.5 – in this case it can be shown that if one randomly selects a copy of gene A and a copy of gene B, then the proteins produced by those copies will have the same ancestry 50% of the time. Thus, these edge weights are the probabilities that compatible proteins interact, assuming that both ancestry types at each locus produce the same amount of protein and the probability that a given protein is involved in a particular interaction does not depend on its ancestry. We then averaged these weights across individuals and across edges in the PPI network to obtain a test statistic. Using this test, we failed to obtain a significant result in either population (*p* = 0.178, CEU, *p* = 0.657, CHBS, one-sided permutation test), which again provides evidence against the DMI hypothesis.

Taken as a whole, we find that while there has been selection against Neanderthal introgression, for example at highly conserved loci, it seems that the negative selection is not due DMIs, which lends more credence to the mutational load hypothesis.

We also note that in contrast to the broad findings presented here, a small number of specific loci have been found to have experienced positive selection for archaic introgression (Racimo et al., 2015; Sams et al., 2016; Racimo et al., 2017).

### 3.4 Patterns of introgression on the X-chromosome

Similar to the map reported by Sankararaman et al. (2014), the introgression map inferred by diCal-admix shows a substantial reduction in Neanderthal introgression on the X-chromosome, when compared to the average introgression on the autosomes. In the results obtained using `diCal-admix`, we observe a five-fold decrease, whereas Sankararaman et al. (2014) reported a six-fold decrease, see Figure 3(a) and Figure 3(b). Similar findings have been reported for Denisovan introgression by Sankararaman et al. (2016). As mentioned in Section 3.3, the authors suggest that this lack of introgression on the X-chromosome might be the result of DMIs causing decreased fertility in male hybrids. Juric et al. (2016) applied a modified version of their inference procedure to obtain estimates for the strength of selection on the X-chromosome as well. Their model only assumes global selection against exonic Neanderthal variation. The confidence intervals obtained by the authors overlap with the confidence intervals obtained for the autosomes and include zero in some cases, but the point estimates provide weak evidence that selection was slightly stronger on the X-chromosome. This suggests that the patterns of introgression on the X-chromosome can, similar to the autosomes, be explained by increased mutational load in Neanderthals without appealing to DMIs.

To explore whether selection against introgressed Neanderthal variation differed between autosomes and the X-chromosome, we performed forward simulations in a Wright-Fisher model, focusing on the dynamics of Neanderthal alleles in the modern human population after the introgression event 1, 900 generations ago. For the autosomes, we modeled each diploid individual to be comprised of two chromosomes. Each chromosome consisted of 5,000 loci, and recombination could act between these loci. The recombination rate was calibrated such that this corresponds to a 150 Mbp chromosome with a recombination rate of 1 .25 × 10^−8^ per generation per base-pair. Similar to Harris and Nielsen (2016), the population size was set to *N* = 1, 860 for the first 900 generations, followed by an instantaneous decrease to *N* = 1,032 with subsequent exponential growth at 0.38% per generation. For computational reasons, we limited the population size to *N* = 10,000, which is reached roughly 300 generations before present. In the generation immediately after the introgression event, the chromosomes of 97% of the individuals in the population carry modern human alleles at all 5,000 loci on both chromosomes, and 3% carry Neanderthal alleles, representing the introgressed individuals. In the subsequent generations, the fitness of an individual is (1 – *s*)^*D*^, with selection coefficient *s*, and *D* denoting the number of Neanderthal alleles that a diploid individual carries. Figure 8(a) depicts the amount of Neanderthal introgression measured in the autosomes in the CEU and the CHB+CHS population, as well as the amount of Neanderthal introgression in a sample of individuals at present from populations simulated with different values for s, repeated 16 times. These simulations suggest that a selection coefficient on the order of *s* = –2 × 10^−5^ is sufficient to explain the observed reduction in Neanderthal ancestry from the initial proportion of 3% on the autosomes. These results are largely consistent with the estimates of selection against introgression provided in (Juric et al., 2016, Table 1), where –3 × 10^−8^ is estimated for the effective selection strength per exonic site. In our simulations, each locus corresponds to 30,000 sites. According to the annotation from the UCSC genome browser (https://genome.ucsc.edu/), the genome-wide density of exonic sites is 2.8%, and thus a simulated locus contains roughly 840 exonic sites. The simulated selection strength of *s* = –2 × 10^−5^ then corresponds to a selection strength of –2.4 × 10^−8^ per exonic site.

We also performed simulations for the X-chromosome to see if the strength of selection is similar on the autosomes and X-chromosome. In our simulation, the fitness of males was determined solely by their single X-chromosome and their Y-chromsome was modeled as selectively neutral. In females, to model X-inactivation, we chose one chromosome randomly to determine the fitness. There are several methods to calibrate the selection coefficient, *s*. Here we chose to calibrate *s* such that, in both females and males, carrying only Neanderthal variants at a certain locus has the same affect on fitness for the X-chromosome as for an autosome. Consequently, to determine the exponent for the fitness in females, the number of Neanderthal alleles on the active chromosome is multiplied by two. In males, the number of Neanderthal alleles on the single X-chromosome is also multiplied by two to obtain the exponent. Note that Juric et al. (2016) used a different calibration, where selection in males is half as strong. Figure 8(b) shows the Neanderthal proportions on the X-chromosome in both populations, and the results of the simulations for different values of *s*. We observe that using this calibration for the selection coefficient, for the same strength of selection, the amount of Neanderthal ancestry in the population at present is reduced on the X-chromosome compared to the autosomes. While this might seem at odds with the reduced effective population size for the X-chromosome, and hence a lower efficacy of selection, it can be explained by the fact that genetic variants in males have a stronger impact than they would in a diploid autosomal population of reduced effective size. Moreover, we observe in Figure8(b) that a selection coefficient not much stronger than *s* = –2 × 10^−5^ can result in the reduction of Neanderthal introgression observed on the X-chromosomes in the 1000 Genomes data.

**Figure 8:**
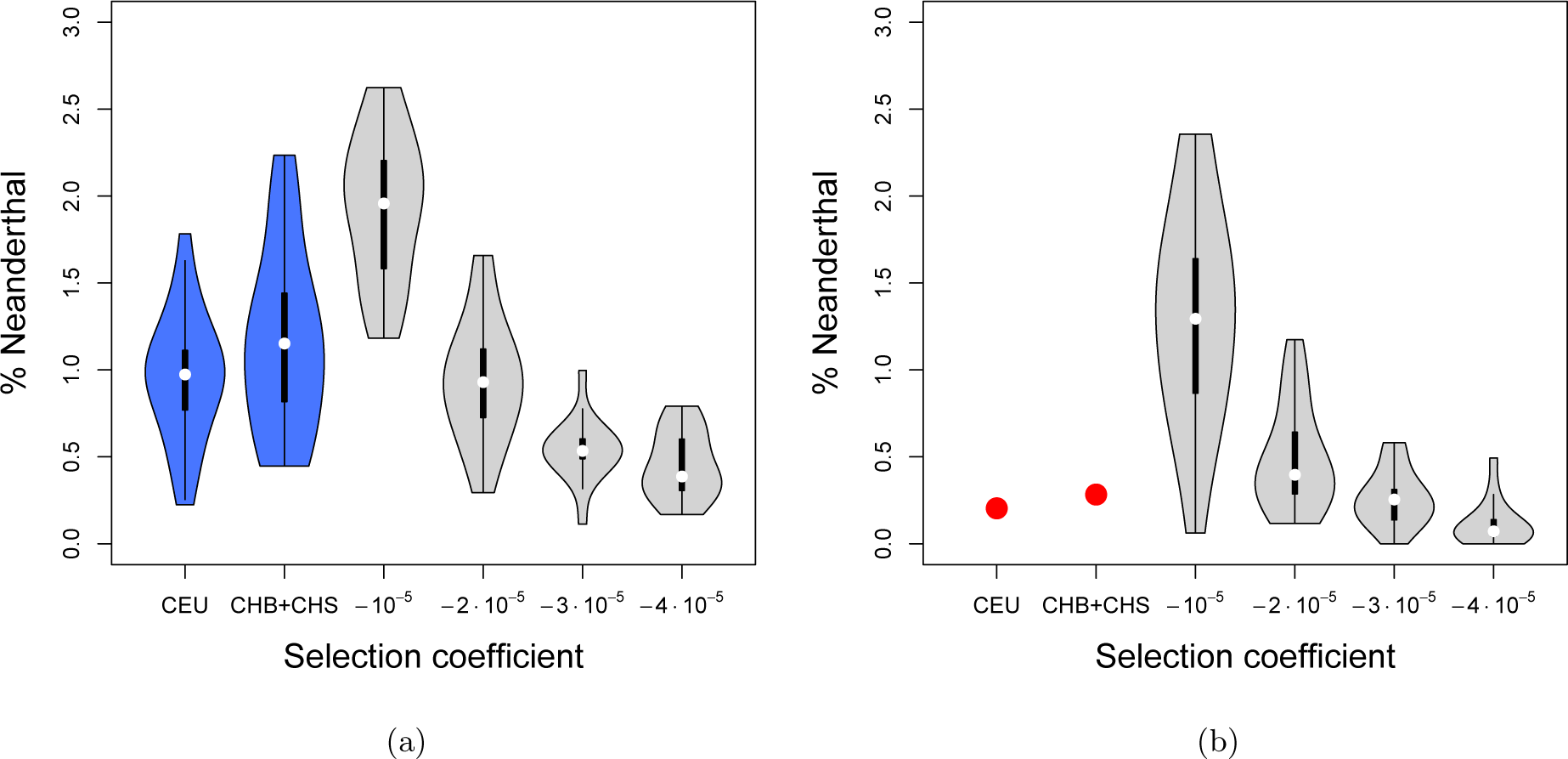
Distribution of amount of Neanderthal introgression on the different chromosomes. Given empirically in CEU and CHB+CHS, and sampled from the present day generation under the Wright-Fisher simulations for different selection coefficients, in 16 replicates. (a) The distributions on the autosomes. (b) The distributions on the X-chromosomes.

To explore the possibility of hybrid male infertility, we modified the simulations for the X-chromosome as follows. In addition to the global selection with coefficient s against Neanderthal alleles at all loci, we designated 0.5% of the 5000 loci to be *incompatibility*-loci. These *incompatibility* loci only affect fitness in male individuals. The fitness is multiplied by 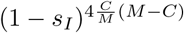, where *s_I_* is the selection coefficient against incompatibility, *M* is the total number of *incompatibility* loci and *C* is the number of Neanderthal alleles a male individual carries at these loci. The exponent is proportional to the number of incompatible pairs. Thus, if an individual carries only modern human or only Neanderthal alleles at these loci, the exponent is zero, and the fitness is not affected. The exponent equals its maximal value of 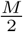 when 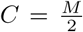, that is, half of the *incompatibility* loci carry the Neanderthal allele, and the other half carries the human allele. Figure 9 depicts the results of the simulations for *s* = –2 × 10^−5^, and different *incompatibility* selection coefficients *s_I_*. Note that the order of magnitude of *s_I_* is higher then *s*, because it is acting on fewer loci. The simulations show that a mechanism like this type of hybrid incompatibility could indeed decrease the introgression further than global weak selection against Neanderthal variants by itself, although such an explanation is not necessary to fit the observed levels of introgression.

**Figure 9:**
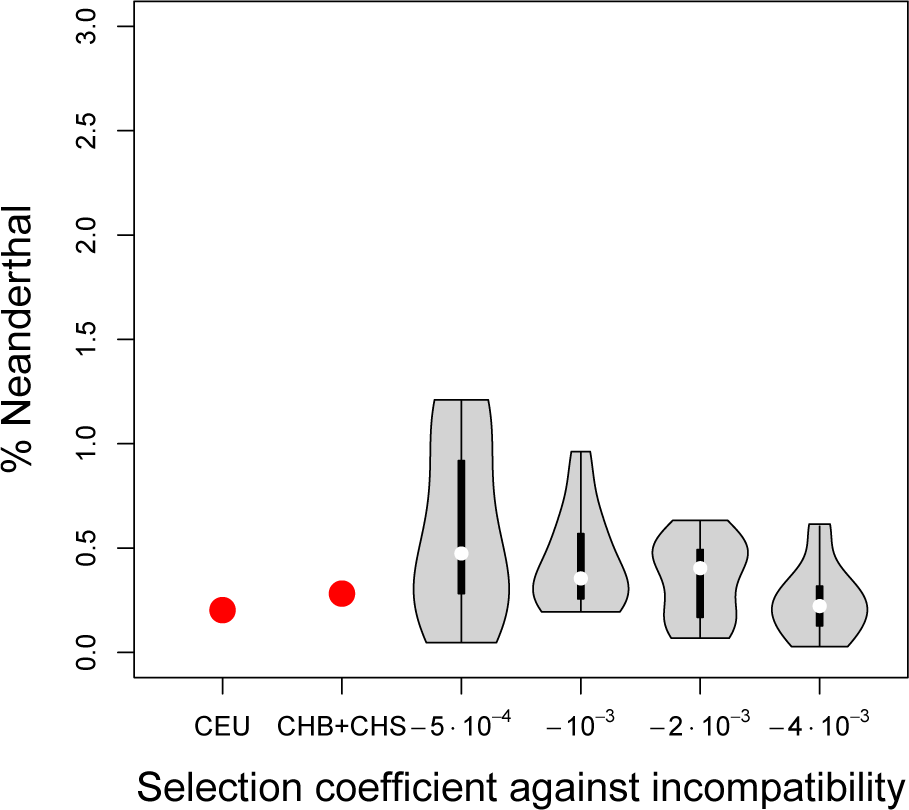
Distribution of amount of Neanderthal introgression on the X-chromosome. Given empirically in CEU and CHB+CHS, and sampled from the present day generation under the Wright-Fisher simulations for different models of male hybrid infertility, in 16 replicates.

The X-chromosome may play an important role for the dynamics of the genetic material that introgressed from Neanderthals into modern humans. We exhibited some of the arguments that have been made to elucidate this role, and performed a simulation study aimed at further understanding the dynamics. However, the evidence that has been collected to date does not seem to be sufficient to fully characterize the importance, and we will list some directions for future research in Section 4.

## 4 Discussion

In this paper, we introduced a modification of the method `diCal 2.0`, which was developed by Steinrücken et al. (2015) to infer complex demographic histories from full-genomic sequence data. We applied this modification (`diCal-admix`) to detect tracts of genetic material in modern non-African individuals that introgressed into the population when non-Africans and Neanderthals exchanged genetic material about 1, 900 generations ago. The method explicitly accounts for the complex underlying demographic history relating these populations. We demonstrated in an extensive simulation study that `diCal-admix` can accurately and efficiently detect tracts of Neanderthal introgression. Furthermore, we applied `diCal-admix` to detect introgression in the individuals sampled from the CEU, CHB, and CHS populations as part of the 1000 Genomes Project (The 1000 Genomes Project Consortium, 2012). We used the YRI individuals as a introgression free reference population, and the Altai Neanderthal (Prüfer et al., 2014) as a reference for the genetic variation in the Neanderthals. We exhibited some of the methodological and empirical differences between `diCal-admix` and previous results reported by Sankararaman et al. (2014) and by Vernot and Akey (2014). While they are generally in good agreement, we observed some differences. This highlights the importance of the development of different methodologies to generate a consensus. We also reported some of the functional implications of introgression, which confirms previously reported findings of wide-spread selection against introgression, enrichment of Neanderthal introgression in certain classes of genes, but a general signal of depleted Neanderthal introgression in conserved regions of the genome. Further, we found some evidence against the hypothesis that this selection is driven by Dobzhansky-Müller incompatibilies, thus lending more credence to mutational load-based hypotheses.

However, the role of the X-chromosome remains intriguing. As in previous studies, we observe a substantially lower amount of Neanderthal introgression on the X-chromosome compared to the autosomes. Sankararaman et al. (2014) hypothesized that this reduction is due to DMIs reducing male fertility, further supported by significantly reduced Neanderthal introgression in genes expressed in testes. However, the authors point out that in other species, such incompatibilities evolve over substantially longer evolutionary times than the divergence between modern humans and Neanderthals. Moreover, the results of our GO-term enrichment analyses did not result in any genes related to testes or infertility. Additionally, other studies (Juric et al., 2016; Harris and Nielsen, 2016) and our simulations suggest that only a moderate strength of selection is required to explain the observed reduction on the X-chromosome. Resolving these questions will require a more comprehensive analysis of larger samples of contemporary genetic data like the Simons Genome Diversity Project (Mallick et al., 2016) and the individuals from Phase III of the 1000 Genomes Project (The 1000 Genomes Project Consortium, 2012). Moreover, additional high-quality data for hominin sister groups (Meyer et al., 2012; Prüfer et al., 2017) will improve the detection of introgression. Detecting introgression in genetic samples of archaic humans (Mathieson et al., 2015) will also allow resolution of the evolutionary trajectory of introgressed genetic material over time. Incorporating the distribution of tracts on an individual level and different modes of selection into the inference frameworks will improve the inference of the strength of selection and allow reliable testing of different models. Additionally, a better understanding of the evolution of incompatibilities, and more careful investigation of the gene content on the X-chromosome will help shed light on the role of the X-chromosome in the Neanderthal introgression landscape. In general, maps of introgressed Neanderthal and Denisovan ancestry will facilitate the interpretation of patterns of human genomic variation and further the understanding of how archaic introgression influenced the trajectory of human evolution.

## Acknowledgements

We thank Sriram Sankararaman for providing us with his introgression calls, and the Rivas lab for making the Global Biobank Engine resource available. This research is supported in part by a National Institutes of Health grant R01-GM094402, and a Packard Fellowship for Science and Engineering. Y.S.S. is a Chan Zuckerberg Biohub investigator.

## Data Accessibility

The marginal posterior probabilities of Neanderthal introgression for the CEU, CHB, and CHS haplotypes computed using `diCal-admix` are available at http://dical-admix.sourceforge.net/

## Supporting Information

Additional supporting information may be found in the online version of this article.

Gene Ontology Analysis Results, ROC and Precision-Recall Curves from Simulated Data, Fine-scale Population Average Introgression

## References

Brandvain, Y., Kenney, A. M., Flagel, L., Coop, G., and Sweigart, A. L. (2014). Speciation and introgression between mimulus nasutus and mimulus guttatus. PLOS Genetics, 10,(6) 1–15.

Brideau, N. J., Flores, H. A., Wang, J., Maheshwari, S., Wang, X., and Barbash, D. A. (2006). Two dobzhansky-muller genes interact to cause hybrid lethality in drosophila. Science, 314,(5803) 1292–1295.

Dannemann, M., Prüfer, K., and Kelso, J. (2017). Functional implications of neandertal introgression in modern humans. Genome Biology, 18,(1) 61.

Dobzhansky, T. (1936). Studies on hybrid sterility. ii. localization of sterility factors in drosophila pseudoobscura hybrids. Genetics, 21,(2) 113–135.

Eden, E., Lipson, D., Yogev, S., and Yakhini, Z. (2007). Discovering motifs in ranked lists of dna sequences. PLOS Computational Biology, 3,(3) 1–15.

Eden, E., Navon, R., Steinfeld, I., Lipson, D., and Yakhini, Z. (2009). Gorilla: a tool for discovery and visualization of enriched go terms in ranked gene lists. BMC Bioinformatics, 10,(1) 48.

Fishman, L. and Willis, J. H. (2001). Evidence for dobzhansky-muller incompatibilites contributing to the sterility of hybrids between mimulus guttatus and m. nasutus. Evolution, 55,(10) 1932–1942.

Fitzpatrick, B. M. (2008). Dobzhansky–muller model of hybrid dysfunction supported by poor burst-speed performance in hybrid tiger salamanders. Journal of Evolutionary Biology, 21,(1) 342–351.

Fitzpatrick, B. M., Johnson, J. R., Kump, D. K., Smith, J. J., Voss, S. R., and Shaffer, H. B. (2010). Rapid spread of invasive genes into a threatened native species. Proceedings of the National Academy of Sciences, 107,(8) 3606–3610.

Gioutlakis, A., Klapa, M. I., and Moschonas, N. K. (2017). Pickle 2.0: A human protein-protein interaction meta-database employing data integration via genetic information ontology. PLOS ONE, 12,(10) 1–17.

Gittelman, R. M., Schraiber, J. G., Vernot, B., Mikacenic, C., Wurfel, M. M., and Akey, J. M. (2016). Archaic hominin admixture facilitated adaptation to out-of-africa environments. Current Biology, 26,(24) 3375–3382.

Global Biobank Engine, (2017). URL http://gbe.stanford.edu/.

Green, R. E., Krause, J., Briggs, A. W., Maricic, T., Stenzel, U., Kircher, M., Patterson, N., Li, H., Zhai, W., Fritz, M. H.-Y., Hansen, N. F., Durand, E. Y., Malaspinas, A.-S., Jensen, J. D., Marques-Bonet, T., Alkan, C., Prüfer, K., Meyer, M., Burbano, H. A., Good, J. M., Schultz, R., Aximu-Petri, A., Butthof, A., Höber, B., Höffner, B., Siegemund, M., Weihmann, A., Nusbaum, C., Lander, E. S., Russ, C., Novod, N., Affourtit, J., Egholm, M., Verna, C., Rudan, P., Brajkovic, D., Kucan, Ž., Gušic, I., Doronichev, V. B., Golovanova, L. V., Lalueza-Fox, C., de la Rasilla, M., Fortea, J., Rosas, A., Schmitz, R. W., Johnson, P. L. F., Eichler, E. E., Falush, D., Birney, E., Mullikin, J. C., Slatkin, M., Nielsen, R., Kelso, J., Lachmann, M., Reich, D., and Pääbo, S. (2010). A draft sequence of the neandertal genome. Science, 328,(5979) 710–722.

Gronau, I., Hubisz, M. J., Gulko, B., Danko, C. G., and Siepel, A. (2011). Bayesian inference of ancient human demography from individual genome sequences. Nature Genetics, 43, 1031 EP -.

Gutenkunst, R. N., Hernandez, R. D., Williamson, S. H., and Bustamante, C. D. (2009). Inferring the joint demographic history of multiple populations from multidimensional snp frequency data. PLOS Genet., 5, (10) e1000695.

Harris, K. and Nielsen, R. (2016). The genetic cost of neanderthal introgression. Genetics, 203,(2) 881–891.

Juric, I., Aeschbacher, S., and Coop, G. (2016). The strength of selection against neanderthal introgression. PLOS Genet., 12,(11) e1006340.

Kelleher, J., Etheridge, A. M., and McVean, G. (2016). Efficient coalescent simulation and genealogical analysis for large sample sizes. PLOS Comput. Biol., 12,(5) e1004842.

Klapa, M. I., Tsafou, K., Theodoridis, E., Tsakalidis, A., and Moschonas, N. K. (2013). Reconstruction of the experimentally supported human protein interactome: what can we learn? BMC Systems Biology, 7, (1) 96.

Kong, A., Thorleifsson, G., Gudbjartsson, D. F., Masson, G., Sigurdsson, A., Jonasdottir, A., Walters, G. B., Jonasdottir, A., Gylfason, A., Kristinsson, K. T., Gudjonsson, S. A., Frigge, M. L., Helgason, A., Thorsteinsdottir, U., and Stefansson, K. (2010). Fine-scale recombination rate differences between sexes, populations and individuals. Nature, 467,(7319) 1099–1103.

Li, N. and Stephens, M. (2003). Modeling linkage disequilibrium and identifying recombination hotspots using single-nucleotide polymorphism data. Genetics, 165,(4) 2213–2233.

Li, Y.-J., Satta, Y., and Takahata, N. (1999). Paleo-demography of the *Drosophila melanogaster* subgroup: application of the maximum likelihood method. Genes & Genetic Systems, 74,(4) 117–127.

Mallick, S., Li, H., Lipson, M., Mathieson, I., Gymrek, M., Racimo, F., Zhao, M., Chennagiri, N., Nordenfelt, S., Tandon, A., Skoglund, P., Lazaridis, I., Sankararaman, S., Fu, Q., Rohland, N., Renaud, G., Erlich, Y., Willems, T., Gallo, C., Spence, J. P., Song, Y. S., Poletti, G., Balloux, F., van Driem, G., de Knijff, P., Romero, I. G., Jha, A. R., Behar, D. M., Bravi, C. M., Capelli, C., Hervig, T., Moreno-Estrada, A., Posukh, O. L., Balanovska, E., Balanovsky, O., Karachanak-Yankova, S., Sahakyan, H., Toncheva, D., Yepiskoposyan, L., Tyler-Smith, C., Xue, Y., Abdullah, M. S., Ruiz-Linares, A., Beall, C. M., Di Rienzo, A., Jeong, C., Starikovskaya, E. B., Metspalu, E., Parik, J., Villems, R., Henn, B. M., Hodoglugil, U., Mahley, R., Sajantila, A., Stamatoyannopoulos, G., Wee, J. T. S., Khusainova, R., Khusnutdinova, E., Litvinov, S., Ayodo, G., Comas, D., Hammer, M. F., Kivisild, T., Klitz, W., Winkler, C. A., Labuda, D., Bamshad, M., Jorde, L. B., Tishkoff, S. A., Watkins, W. S., Metspalu, M., Dryomov, S., Sukernik, R., Singh, L., Thangaraj, K., Pääbo, S., Kelso, J., Patterson, N., and Reich, D. (2016). The simons genome diversity project: 300 genomes from 142 diverse populations. Nature, 538,(7624) 201–206.

Malnic, B., Godfrey, P. A., and Buck, L. B. (2004). The human olfactory receptor gene family. Proceedings of the National Academy of Sciences of the United States of America, 101,(8) 2584–2589.

Mathieson, I., Lazaridis, I., Rohland, N., Mallick, S., Patterson, N., Roodenberg, S. A., Harney, E., Stewardson, K., Fernandes, D., Novak, M., Sirak, K., Gamba, C., Jones, E. R., Llamas, B., Dryomov, S., Pickrell, J., Arsuaga, J. L., de Castro, J. B., Carbonell, E., Gerritsen, F., Khokhlov, A., Kuznetsov, P., Lozano, M., Meller, H., Mochalov, O., Moiseyev, V., Guerra, M. A. R., Roodenberg, J., Vergès, J. M., Krause, J., Cooper, A., Alt, K. W., Brown, D., Anthony, D., Lalueza-Fox, C., Haak, W., Pinhasi, R., and Reich, D. (2015). Genome-wide patterns of selection in 230 ancient eurasians. Nature, 528,(7583) 499–503.

Meyer, M., Kircher, M., Gansauge, M.-T., Li, H., Racimo, F., Mallick, S., Schraiber, J. G., Jay, F., Prüfer, K., de Filippo, C., Sudmant, P. H., Alkan, C., Fu, Q., Do, R., Rohland, N., Tandon, A., Siebauer, M., Green, R. E., Bryc, K., Briggs, A. W., Stenzel, U., Dabney, J., Shendure, J., Kitzman, J., Hammer, M. F., Shunkov, M. V., Derevianko, A. P., Patterson, N., Andrés, A. M., Eichler, E. E., Slatkin, M., Reich, D., Kelso, J., and Pääbo, S. (2012). A high-coverage genome sequence from an archaic denisovan individual. Science, 338,(6104) 222–226.

Myers, S., Bottolo, L., Freeman, C., McVean, G., and Donnelly, P. (2005). A fine-scale map of recombination rates and hotspots across the human genome. Science, 310,(5746) 321–324.

O’Leary, N. A., Wright, M. W., Brister, J. R., Ciufo, S., Haddad, D., McVeigh, R., Rajput, B., Robbertse, B., Smith-White, B., Ako-Adjei, D., Astashyn, A., Badretdin, A., Bao, Y., Blinkova, O., Brover, V., Chetvernin, V., Choi, J., Cox, E., Ermolaeva, O., Farrell, C. M., Goldfarb, T., Gupta, T., Haft, D., Hatcher, E., Hlavina, W., Joardar, V. S., Kodali, V. K., Li, W., Maglott, D., Masterson, P., McGarvey, K. M., Murphy, M. R., O’Neill, K., Pujar, S., Rangwala, S. H., Rausch, D., Riddick, L. D., Schoch, C., Shkeda, A., Storz, S. S., Sun, H., Thibaud-Nissen, F., Tolstoy, I., Tully, R. E., Vatsan, A. R., Wallin, C., Webb, D., Wu, W., Landrum, M. J., Kimchi, A., Tatusova, T., DiCuccio, M., Kitts, P., Murphy, T. D., and Pruitt, K. D. (2016). Reference sequence (refseq) database at ncbi: current status, taxonomic expansion, and functional annotation. Nucleic Acids Research, 44,(D1) D733–D745.

Orr, H. A. (1995). The population genetics of speciation: the evolution of hybrid incompatibilities. Genetics, 139,(4) 1805–1813.

Paten, B., Herrero, J., Beal, K., Fitzgerald, S., and Birney, E. (2008). Enredo and pecan: Genome-wide mammalian consistency-based multiple alignment with paralogs. Genome Res., 18,(11) 1814–1828.

Paul, J. S. and Song, Y. S. (2010). A principled approach to deriving approximate conditional sampling distributions in population genetics models with recombination. Genetics, 186,(1) 321–338.

Paul, J. S., Steinrücken, M., and Song, Y. S. (2011). An accurate sequentially markov conditional sampling distribution for the coalescent with recombination. Genetics, 187,(4) 1115–1128.

Plagnol, V. and Wall, J. D. (2006). Possible ancestral structure in human populations. PLOS Genet., 2,(7) e105.

Presgraves, D. C. (2008). Sex chromosomes and speciation in drosophila. Trends Genet., 24,(7) 336–343.

Prufer, K., Racimo, F., Patterson, N., Jay, F., Sankararaman, S., Sawyer, S., Heinze, A., Renaud, G., Sudmant, P. H., de Filippo, C., Li, H., Mallick, S., Dannemann, M., Fu, Q., Kircher, M., Kuhlwilm, M., Lachmann, M., Meyer, M., Ongyerth, M., Siebauer, M. F., Theunert, C., Tandon, A., Moorjani, P., Pickrell, J., Mullikin, J. C., Vohr, S. H., Green, R. E., Hellmann, I., Johnson, P. L. F., Blanche, H., Cann, H., Kitzman, J. O., Shendure, J., Eichler, E. E., Lein, E. S., Bakken, T. E., Golovanova, L. V., Doronichev, V. B., Shunkov, M. V., Derevianko, A. P., Viola, B., Slatkin, M., Reich, D., Kelso, J., and Pääbo, S. (2014). The complete genome sequence of a Neanderthal from the Altai Mountains. Nature, 505,(7481) 43–49.

Prufer, K., de Filippo, C., Grote, S., Mafessoni, F., Korlević, P., Hajdinjak, M., Vernot, B., Skov, L., Hsieh, P., Peyrégne, S., Reher, D., Hopfe, C., Nagel, S., Maricic, T., Fu, Q., Theunert, C., Rogers, R., Skoglund, P., Chintalapati, M., Dannemann, M., Nelson, B. J., Key, F. M., Rudan, P., Kućan, Ž., Gušić, I., Golovanova, L. V., Doronichev, V. B., Patterson, N., Reich, D., Eichler, E. E., Slatkin, M., Schierup, M. H., Andrés, A., Kelso, J., Meyer, M., and Püübo, S. (2017). A high-coverage neandertal genome from vindija cave in croatia. Science. aao1887.

Racimo, F., Sankararaman, S., Nielsen, R., and Huerta-Sánchez, E. (2015). Evidence for archaic adaptive introgression in humans. Nature Reviews Genetics, 16, 359 EP -.

Racimo, F., Marnetto, D., and Huerta-Sánchez, E. (2017). Signatures of archaic adaptive introgression in present-day human populations. Molecular Biology and Evolution, 34,(2) 296–317.

Rogers, R. L. (2015). Chromosomal rearrangements as barriers to genetic homogenization between archaic and modern humans. Molecular Biology and Evolution, 32,(12) 3064–3078.

Sams, A. J., Dumaine, A., Nedéléc, Y., Yotova, V., Alfieri, C., Tanner, J. E., Messer, P. W., and Barreiro, L. B. (2016). Adaptively introgressed neandertal haplotype at the oas locus functionally impacts innate immune responses in humans. Genome Biology, 17,(1) 246.

Sankararaman, S., Patterson, N., Li, H., Pääbo, S., and Reich, D. (2012). The date of interbreeding between neandertals and modern humans. PLOS Genet., 8,(10) e1002947.

Sankararaman, S., Mallick, S., Dannemann, M., Prufer, K., Kelso, J., Paabo, S., Patterson, N., and Reich, D. (2014). The genomic landscape of neanderthal ancestry in present-day humans. Nature, 507,(7492) 354–357.

Sankararaman, S., Mallick, S., Patterson, N., and Reich, D. (2016). The combined landscape of denisovan and neanderthal ancestry in present-day humans. Curr. Biol., 26,(9) 1241–1247.

Scally, A. and Durbin, R. (2012). Revising the human mutation rate: implications for understanding human evolution. Nat Rev Genet, 13,(10) 745–753.

Schumer, M., Xu, C., Powell, D., Durvasula, A., Skov, L., Holland, C., Sankararaman, S., Andolfatto, P., Rosenthal, G., and Przeworski, M. (2017). Natural selection interacts with the local recombination rate to shape the evolution of hybrid genomes. bioRxiv.

Shaffer, H. B. and McKnight, M. L. (1996). The polytypic species revisited: Genetic differentiation and molecular phylogenetics of the tiger salamander ambystoma tigrinum (amphibia: Caudata) complex. Evolution, 50,(1) 417–433.

Siepel, A., Bejerano, G., Pedersen, J. S., Hinrichs, A. S., Hou, M., Rosenbloom, K., Clawson, H., Spieth, J., Hillier, L. W., Richards, S., Weinstock, G. M., Wilson, R. K., Gibbs, R. A., Kent, W. J., Miller, W., and Haussler, D. (2005). Evolutionarily conserved elements in vertebrate, insect, worm, and yeast genomes. Genome Research, 15,(8) 1034–1050.

Simonti, C. N., Vernot, B., Bastarache, L., Bottinger, E., Carrell, D. S., Chisholm, R. L., Crosslin, D. R., Hebbring, S. J., Jarvik, G. P., Kullo, I. J., Li, R., Pathak, J., Ritchie, M. D., Roden, D. M., Verma, S. S., Tromp, G., Prato, J. D., Bush, W. S., Akey, J. M., Denny, J. C., and Capra, J. A. (2016). The phenotypic legacy of admixture between modern humans and neandertals. Science, 351,(6274) 737–741.

Slatkin, M. and Racimo, F. (2016). Ancient dna and human history. Proceedings of the National Academy of Sciences, 113,(23) 6380–6387.

Steinrücken, M., Kamm, J.,, and Song, Y. (2015). Inference of complex population histories using whole-genome sequences from multiple populations. Preprint at: http://dx.doi.org/10.1101/026591.

Steinrücken, M., Paul, J. S., and Song, Y. S. (2013). A sequentially Markov conditional sampling distribution for structured populations with migration and recombination. Theor. Popul. Biol., 87, 51–61.

Sudlow, C., Gallacher, J., Allen, N., Beral, V., Burton, P., Danesh, J., Downey, P., Elliott, P., Green, J., Landray, M., Liu, B., Matthews, P., Ong, G., Pell, J., Silman, A., Young, A., Sprosen, T., Peakman, T., and Collins, R. (2015). Uk biobank: An open access resource for identifying the causes of a wide range of complex diseases of middle and old age. PLOS Medicine, 12,(3) 1–10.

The 1000 Genomes Project Consortium. (2012). An integrated map of genetic variation from 1,092 human genomes. Nature, 491,(7422) 56–65.

Veeramah, K. R., Wegmann, D., Woerner, A., Mendez, F. L., Watkins, J. C., Destro-Bisol, G., Soodyall, H., Louie, L., and Hammer, M. F. (2012). An early divergence of khoesan ancestors from those of other modern humans is supported by an abc-based analysis of autosomal resequencing data. Molecular Biology and Evolution, 29,(2) 617–630.

Vernot, B. and Akey, J. M. (2014). Resurrecting surviving neandertal lineages from modern human genomes. Science, 343,(6174) 1017–1021.

Vernot, B. and Akey, J. M. (2015). Complex history of admixture between modern humans and neandertals. The American Journal of Human Genetics, 96,(3) 448 - 453.

Vernot, B., Tucci, S., Kelso, J., Schraiber, J. G., Wolf, A. B., Gittelman, R. M., Dannemann, M., Grote, S., McCoy, R. C., Norton, H., Scheinfeldt, L. B., Merriwether, D. A., Koki, G., Friedlaender, J. S., Wakefield, J., Pääbo, S., and Akey, J. M. (2016). Excavating neandertal and denisovan dna from the genomes of melanesian individuals. Science aad9416.

